# Resolvin D2-GPR18 Signaling on Myeloid Cells Limits Plaque Necrosis

**DOI:** 10.1101/2023.04.03.535493

**Authors:** Masharh Lipscomb, Sean Walis, Michael Marinello, Hebe Agustina Mena, Matthew Spite, Gabrielle Fredman

**Author notes:** Correspondence should be addressed to G.F., The Department of Molecular and Cellular Physiology, Albany Medical College, Albany Medical College, Albany, NY 12208, USA.

## Abstract

**Introduction/Objective:** Dysregulated inflammation-resolution programs are associated with atherosclerosis progression. Inflammation-resolution is in part mediated by Resolvins, including Resolvin D2 (RvD2). RvD2, which activates a G-protein coupled receptor (GPCR) called GPR18, limits plaque progression. Cellular targets of RvD2 are not known.

**Approach and Results:** Here we developed humanized GPR18 floxed (“fl/fl”) and a myeloid (Lysozyme M Cre) GPR18 knockout (mKO) mouse. We functionally validated this model by assessing efferocytosis in bone marrow derived macrophages (BMDMs) and found that RvD2 enhanced efferocytosis in the fl/fl, but not in the mKO BMDMs. We employed two different models to evaluate the role of GPR18 in atherosclerosis. We first used the PCSK9-gain of function approach and found increased necrosis in the plaques of the mKO mice compared with fl/fl mice. Next, we performed a bone marrow transfer of fl/fl or mKO bone marrow into *Ldlr*^-/-^ recipients. For these experiments, we treated each genotype with either Veh or RvD2 (25 ng/mouse, 3 times/week for 3 weeks). Myeloid loss of GPR18 resulted in significantly more necrosis and cleaved caspase-3^+^ cells compared with fl/fl transplanted mice. RvD2 treatment decreased plaques necrosis and cleaved caspase-3^+^ cells in fl/fl, but not in the mKO transplanted mice.

**Conclusions:** These results are the first to suggest a causative role for endogenous RvD2 signaling on myeloid cells in limiting plaque necrosis. Moreover, these results provide a mechanistic basis for RvD2 as a therapy limiting plaque necrosis.

## Introduction

Non-resolving inflammation is emerging as a major underpinning in the progression of atherosclerosis (1, 2). Therapeutically targeting inflammation is difficult because of unintended consequences to host defense mechanisms which include an increased susceptibility to infection. Therefore, there is a need to understand how to control inflammation without compromising host defense programs. Resolution is an active process that is regulated in part through the biosynthesis of specific lipid mediators called specialized pro-resolving mediators (SPMs), including Resolvins (3). Resolvin D2 (RvD2), for example, is a potent ligand that binds to a G-protein coupled receptor called GPR18 (4). RvD2 acts on macrophages and enhances phagocytosis and efferocytosis in a GPR18 dependent manner (4). An important function of RvD2 is that it controls infection in mice by stimulating host defense mechanisms, like neutrophil containment and killing of bacteria (5). Therefore, RvD2 is not immunosuppressive in mice and may be an important therapeutic in treating long-term progressive diseases like atherosclerosis.

Along these lines, RvD2 treatment to Apoe^-/-^ mice limits plaque necrosis in mice and increases the expression of pro-reparative markers on macrophages (6, 7). SPMs, including RvD2 were also shown to decrease as atherosclerosis progressed which suggested an association between lowered levels of plaque SPMs and atherosclerosis progression (6, 8). However, major gaps remain in our understanding of RvD2 signaling in atherosclerosis. For example, causation experiments for the endogenous role of RvD2 signaling in atherosclerosis as well as which cellular target(s) RvD2 acts upon to limit plaque necrosis are not known.

To address this gap, we generated a new humanized GPR18 floxed mouse (fl/fl) to test if loss of GPR18 on myeloid cells impacts plaque progression. Briefly, we found that GPR18 is regulated by atherogenic stimuli and that the loss of GPR18 on myeloid cells drives plaque necrosis. We also found that RvD2’s ability to limit plaque necrosis was in part due to GPR18 signaling on myeloid cells. These data suggest a causative role for endogenous RvD2 signaling and provide the first evidence of a myeloid RvD2- GPR18 targeted signaling event to limit necrosis.

## Material and Methods

### Humanized GPR18 floxed mouse

Human GPR18 (hGPR18) containing two loxp sites was inserted following the excision of Murine GPR18. NeoSTOP cassette mice containing two FRT sites were crossed with a FLPe mouse flippase (B6.129S4-441 Gt(ROSA)26Sortm1(FLP1)Dym/RainJ; Stock Number: 009086, JAX). Mice were then subjected to backcrossing by breeding them with C57BL6 mice producing hGPR18 floxed mice (fl/fl). Myeloid specific KO (mKO) mice were produced by crossing fl/fl mice as above with mice containing a LysMCre (The Jackson Laboratory).

### Murine atherosclerosis

All mice were socially housed in standard cages at 22°C under a 12 hr. light and 12 hr. dark cycle at the Albany Medical College animal facility. Also, all procedures were performed according to the animal protocols approved by the Albany Medical College Institutional Animal Care and Use Committee.

### Bone marrow transplantation

Eight-week-old *Ldlr*^-/-^ were purchased (The Jackson Laboratory) and subjected to lethal radiation for complete ablation of bone marrow. Radiation was given in two doses being 4.75 Gy, with 4 h between doses. After the second dose of radiation, bone marrow from fl/fl and mKO mice was intravenously injected into these *Ldlr*^-/-^ mice. The mice were given antibiotic water (SMZ and TMP; catalog no. NDC 65862-496-47; Aurobindo Pharma) and a 6-wk recovery period. After 6 weeks, mice were fed a Western Diet (WD, Envigo TD.88137) for 11 weeks to allow for development of atherosclerosis, after which mice were randomly assigned to receive intraperitoneal injections of either vehicle/PBS or RvD2 (25 ng/mouse) 3 times/week for 3 weeks while still on WD. The total duration of WD feeding was 14 weeks for these experiments.

### Atherosclerotic Lesion Analysis

Aortic roots from vehicle/PBS treated or RvD2 treated mice and fl/fl or mKO mice were harvested for histological analysis. Lesional and necrotic analyses were performed on hematoxylin and eosin (H&E)-stained lesional cross-sections and were quantified using an Olympus camera and Olympus DP2-BSW software. Briefly, frozen specimens were immersed in OCT, cryosectioned, and 10µm sections were placed on glass slides. Atherosclerotic lesion area, defined as the region from the internal elastic lamina to the lumen, was quantified by taking the average of 6 sections spaced around 24µm apart beginning at the base of the aortic root (9).

### Lesional Cleaved Caspase 3 Staining

Frozen aortic root sections were fixed with ice-cold 100% methanol for 10 min. Fixed sections were incubated with blocking buffer (1% Bovine Serum Albumin (BSA) + 10% Goat Serum) for 1 hour at room temperature. Sections were then incubated with rabbit anti-mouse Cleaved Caspase-3 (10) (Cell Signaling Technologies, catalog no. 9978S) at 1:100 dilution overnight at 4°C in 1% BSA. On the following day, sections were then washed 3x with 1x PBS and incubated with Alexa Fluor 647 goat anti-rabbit secondary Ab (Thermo Fisher Scientific, catalog no. A-21245) at 1:200 dilution in 1% BSA + 5% Goat Serum for 2 hrs. at room temperature. Sections were then washed with 1x PBS. Nuclei were stained with Hoechst (Invitrogen, catalog no. H3570) for 10 mins, and images were acquired immediately on a Leica SPE confocal microscope at 40x magnification and analyzed with Image J software. Cleaved Caspase-3 positive cells were counted and expressed as a percentage of total lesional cells.

### Blood Glucose and cholesterol measurements

WD-fed mice were subjected to a 6-hour fast, after which blood glucose was assessed by blood droplet from tail vein with the OneTouch Blood Glucose Meter (catalog no. 021-911). WD-diet fed *Ldlr*^-/-^ mice were sacrificed, and blood was collected in 10% EDTA by retro-orbital bleeding procedure. Immediately after collection, blood was centrifuged at full speed for 30 min at 4°C, and a cholesterol assay (catalog no. 999-02601; Wako Diagnostics) was performed according to manufacturer’s instruction.

### In vitro efferocytosis

Experiments were done as in (9). Briefly, BMDMs from C57BL6, fl/fl and mKO mice were plated on 24-well plates (200,000/cells per well) and after ∼16 hrs. treated with Vehicle or 1 nM RvD2 for 20 mins (37°C, 5% CO_2_) prior to performing efferocytosis. In parallel, Jurkats were enumerated and then stained with PKH26 (Sigma, catalog # PKH26GL) according to the manufacturer’s instructions. Excess dye was removed by washing and the Jurkats were resuspend in RPMI containing 10% FBS. To induce apoptosis, Jurkats were then exposed to UV radiation (0.16 Amps, 115 Volts, 254 nm wavelength) for 15 mins at room temperature and then placed in an incubator (37°C, 5% CO_2_) for 3 hrs. (11). Apoptotic Jurkats were co-cultured with BMDMs in a 3:1 ratio for an additional 1 hr. in 37°C, 5% CO_2_. Excess apoptotic cells were removed by washing ∼3x with PBS. The cells were then immediately fixed with 4% formalin and subjected to fluorescence imaging with a Biorad ZOE Fluorescent Cell Imager. Six-to-seven different fields were acquired per well/group and an efferocytic event was considered a macrophage containing red apoptotic cells. Results were expressed as the percent of efferocytosis per total macrophages.

### Statistical analysis

For all *in vivo* studies, mice were randomly assigned to their respective groups. Results are represented as mean ± SEM. Prism (GraphPad, La Jolla, CA) software was used for statistical analysis, and statistical differences were determined using the two- tailed Student’s t test, Mann Whitney or one-way ANOVA, with multiple comparison post hoc analyses depending on the contexts. Details regarding statistical tests can be found in the figure legends.

## Results

### GPR18 expression is increased by pro-inflammatory and atherogenic stimuli

Because GPR18 is expressed in human plaques (7), we first questioned whether pro-inflammatory stimuli that are also associated with atherosclerosis progression modulate the expression of GPR18. For these experiments, bone marrow derived macrophages (BMDMs) were stimulated with 100 ng/mL of IL-6 or IL-1β for 24 hrs. We observed that *Gpr18* expression was significantly increased by IL-6 and IL-1β (**SFig 1A**) treatment suggesting that inflammation may poise BMDMs to receive GPR18 signals. These data provide rationale for deeper exploration of GPR18 function and cell type specificity in atherosclerosis.

**Fig 1.**
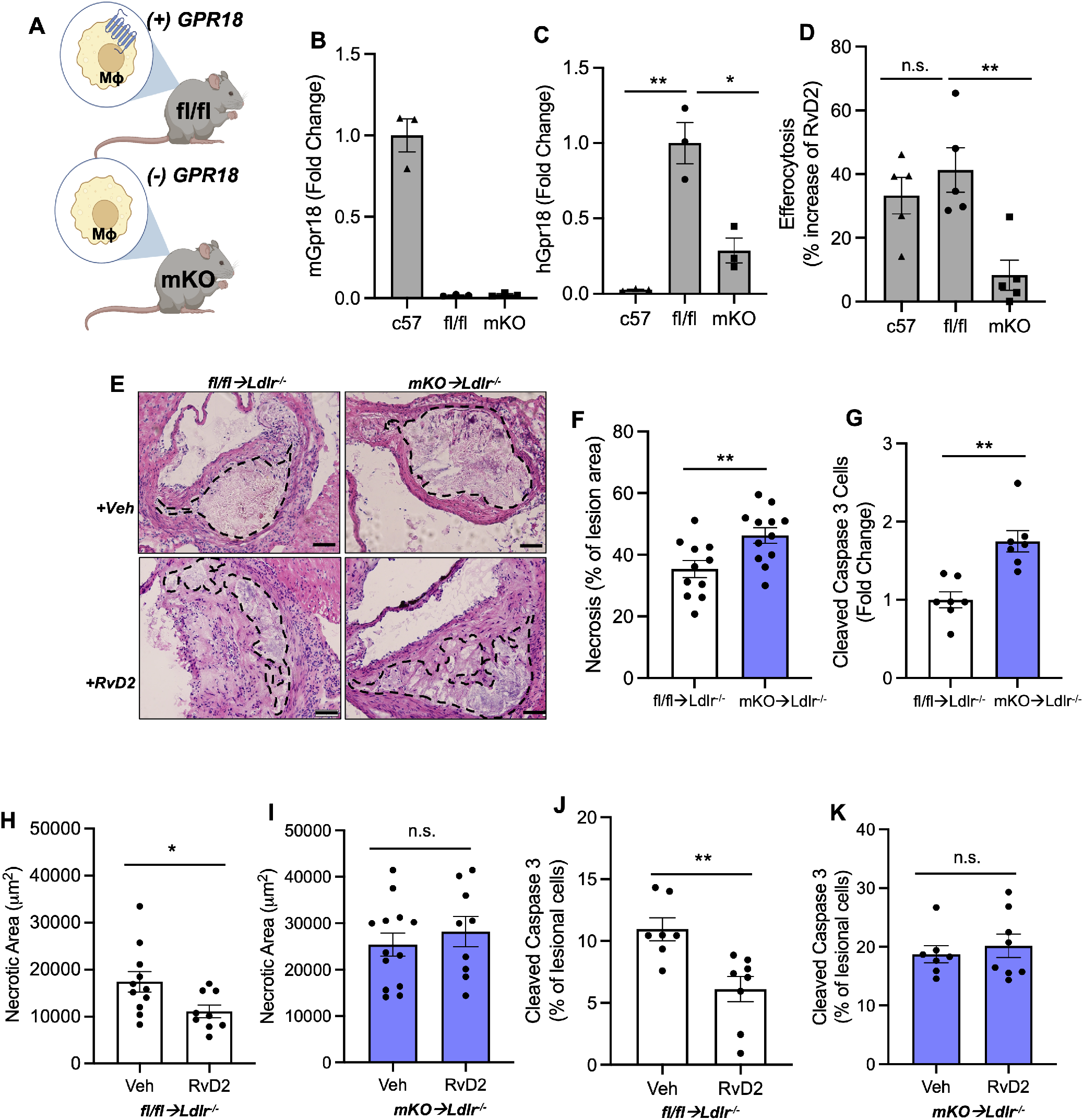
RvD2-Gpr18 signaling on myeloid cell limits plaque necrosis. **(A)** Scheme depicting both fl/fl and mKO humanized GPR18 mice. Quantitative PCR was performed to assess (**B**) murine *Gpr18 and* (**C**) human *GPR18* and expression in C57BL6, fl/fl, and mKO BMDMs. (**D**) Efferocytosis assays were done as described in the methods section. Images were acquired on a Zoe Fluorescence imager and data were analyzed with ImageJ. The data are presented as the percent increase in efferocytosis by RvD2. **(E-G)** Fl/fl or mKO bone marrow was transferred into *Ldlr*^*-/-*^ mice fed a WD for 14 weeks. Aortic root sections were analyzed for (**F**) Lesion and (**G**) necrotic area by serial H&E images and Olympus software. Veh or RvD2 (25 ng/mouse) was given to *Ldlr*^*-/-*^ mice that received bone marrow transfers of fl/fl or mKO cells. Necrotic area is shown in Veh and RvD2 treated **(H)** fl/fl→*Ldlr*^*-/-*^ and (**I**) mKO→*Ldlr*^*-/-*^ mice. Aortic roots were subjected to immunofluorescence staining of cleaved caspase-3. Images were taken with a Leica SPE microscope at 40x magnification and quantified using Image J. The percent cleaved caspase-3^+^ positive cells Veh and RvD2 treated **(J)** fl/fl→*Ldlr*^*-/-*^ and **(K)** mKO→*Ldlr*^*-/-*^. All results are mean ± SEM and each dot represents an individual experiment or mouse. N.s. = not significant *p<0.05, **p<0.01. Panel B-D, Kruskal-Wallis with Dunn’s post-test and Panels F-K are Mann-Whitney test.

### Loss of GPR18 on Myeloid Cells, limits RvD2’s ability to enhance efferocytosis

RvD2 limits atheroprogression in mice (6, 7) but the cell types through which RvD2 acts to limit atherosclerosis are not known. We generated a new mouse in which murine *Gpr18* was replaced with a floxed human *GPR18* allele (**Fig. 1A**). Mice expressing the human floxed *GPR18* gene (referred to as “fl/fl”) were crossed with a *LysM*-driven Cre line to generate myeloid-specific GPR18 knockout mice (or “mKO”). Quantitative RT-PCR was used to determine loss of murine *Gpr18*. Both fl/fl and mKO mice had virtually undetectable murine Gpr18, compared with C57BL6 control mice (**Fig. 1B**). The fl/fl mice had significantly more human GPR18 expression compared with mKO, suggesting that the LysMCre successfully reduced GPR18 expression in the mKO mice (**Fig. 1C**).

Next, we evaluated murine BMDMs to determine whether human GPR18 was functional in murine cells. We performed an *in vitro* efferocytosis assay where BMDMs were treated with Veh or RvD2 (1 nM) for 15 minutes, after which PHK26-labelled apoptotic Jurkats were added in a 1:3 ratio. After 60 minutes, the apoptotic cells were washed off and efferocytosis was assessed by fluorescence imaging and ImageJ analysis. We found that RvD2 enhanced efferocytosis in BMDMs from C57BL6 control mice and the fl/fl mice, suggesting that the human GPR18 was as functional as murine GPR18 in murine cells (**Fig. 1D**). Importantly, RvD2 did not increase efferocytosis in the mKO BMDMs (**Fig. 1D**), demonstrating that we successfully removed both murine and human GPR18 in macrophages and that RvD2’s ability to enhance efferocytosis was dependent on GPR18.

### Loss of Gpr18 on myeloid cells drives plaque necrosis

Because efficient efferocytosis is critical to limit plaque necrosis, we next questioned whether the mKO mice would have worsened necrosis. First, we injected an AAV8 gain of function PCSK9 (AAV8-gof-PCSK9) virus into female fl/fl or mKO mice and fed them a WD for 14 weeks. Aortic roots were harvested, staining with H&E and serial sections were analyzed for lesional and necrotic area. Representative H&E images are shown and depict larger areas of necrosis in the mKO compared with fl/fl (**SFig. 2A**). Indeed, quantification reveals that there was no significant change in lesional area between the groups (**SFig. 2B**). There was a significant increase in necrosis (**SFig. 2C**) in mKO mice compared with fl/fl controls. Importantly, there were no statistical differences on total cholesterol (**SFig. 2C**) and body weight (**SFig. 2D**) between the mKO and fl/fl mice. These data suggest that loss of Gpr18 on myeloid cells increases necrosis and worsens plaque progression.

Next, we performed a bone marrow transfer of male fl/fl or mKO mice into male Ldlr^-/-^. For these experiments, fl/fl-*Ldlr*^-/-^ or mKO-*Ldlr*^-/-^ mice were fed a WD for 11 weeks, followed by Veh or RvD2 (25 ng/mouse, 3x/week) for 3 additional weeks on WD. Aortic roots were harvested, and H&E staining was performed (**Fig. 1E**). First mKO-*Ldlr*^- /-^ mice had increased plaque necrosis, compared with fl/fl-*Ldlr*^-/-^, which is consistent with results in **SFig. 2C**. The lesion area between the fl/fl and mKO groups (49,347µm^2^ ± 4,908 and 54,532µm^2^ ± 4,692 respectively) were not different, which is also consistent with results in **SFig. 2B**. Immunofluorescence staining of cleaved caspase-3 was used to determine the number of apoptotic cells in plaques. Indeed, mKO-*Ldlr*^-/-^ had significantly more cleaved caspase-3^+^ cells, compared with fl/fl-*Ldlr*^-/-^ (**Fig. 1G**). These data suggest that the loss of myeloid GPR18 drives plaque necrosis and the accumulation of apoptotic cells. Collectively these data support the concept that endogenous GPR18 ligands, like RvD2, acts on myeloid cells to limits plaque necrosis.

### RvD2 limits plaque necrosis, in part via GPR18 Signaling on myeloid cells

To determine whether RvD2 engaged myeloid Gpr18 to limit atherosclerosis, we treated fl/fl-*Ldlr*^-/-^ or mKO-*Ldlr*^-/-^ mice as described above. We observed that RvD2 significantly decreased plaque necrosis in fl/fl-*Ldlr*^-/-^ mice (**Fig. 1H**) but not in mKO-*Ldlr*^-/-^ (**Fig. 1I**). Lastly, we also observed a similar pattern in which RvD2 significantly decreased the percent of cleaved caspase3^+^ cells in fl/fl- *Ldlr*^-/-^ mice (**Fig. 1J**) but not in mKO-*Ldlr*^-/-^. (**Fig. 1K**). Importantly, total cholesterol, fasting blood glucose and body weight were not different between the groups (**SFig. 3A-C**). These data suggest that RvD2 limits plaque necrosis and apoptotic cells in part through myeloid GPR18 signaling.

## Discussion

This study provides evidence that the loss of RvD2’s receptor on myeloid cell leads to increased accumulation of dead cells and necrosis in plaques. Also, we demonstrate that RvD2 treatment reduce accumulation of dead cells and plaque necrosis in fl/fl control mice but not the Gpr18 mKO mice. These results, suggest that RvD2, in part acts, through its GPCR to limit death and necrosis in plaques. Overall, we provide evidence that there is an uncoupling of a pro-resolving circuit that may be amenable as a therapeutic intervention.

Atherosclerosis is propagated by chronic non-resolving inflammation and so understanding endogenous circuits that may dysregulate resolution is of interest. Here we provide evidence that RvD2-Gpr18 is an example of a dysregulated inflammation-resolution axis. We and others observed that pro-inflammatory factors increase Gpr18 expression. These data, in the context of the literature in which *Viola et al* observed RvD2 and Maresin 1 levels decreased in plaques as atherosclerosis progresses, suggests an uncoupling of a pro-resolving circuit that is required to limit plaque necrosis. Conceptually, self-limited inflammation jump starts resolution responses (12) and so a break in this inflammation-resolution circuit can, in part, lead to chronicity.

Another recent example of a dysregulated RvD2-GPR18 resolution axis in in human sepsis. Jundi B *et al*, observed that *GPR18* expression is increased in patients with sepsis (13). Moreover, *ex vivo* treatment of peripheral blood myeloid cells, like monocytes and neutrophils with nanomolar concentration of RvD2 partially reversed sepsis-induced changes, which suggests an uncoupling of the GPR18-RvD2 axis in human sepsis.

Aging is a major risk factor for atherosclerosis and Gpr18 expression is increased in bone marrow monocytes from old mice compared with young (14). These data also suggest that aging may uncouple the RvD2-GPR18 axis which warrants further exploration. Along these lines, bone marrow transplanted from old mice has increased atherosclerosis progression compared with bone marrow transplanted from young mice (15). Aging is also associated with impaired macrophage efferocytosis (16), which is an important program that limits plaque necrosis. RvD2 enhances efferocytosis in other contexts and so a deeper exploration of the RvD2-GPR18 axis plays in age-related processes would be of interest to explore in the future. Our work further demonstrates that bone marrow cells and macrophages are critical effectors of plaque atheroprogression (17) and highlights the importance of investigating the role of hematopoietic cells specifically and their role in atherosclerosis.

Previous studies demonstrated the pro-resolving function of the RvD2-GPR18 axis with the use of Gpr18 global knockout mice in the context of bacterial infection and hindlimb ischemia reperfusion (4, 18). Work herein is the first report to suggest the importance of myeloid RvD2-GPR18 signaling in limiting plaque necrosis. These findings are important because they provide further evidence how as to the cellular targets through RvD2 acts and may aid in better targeting a RvD2 treatment. Indeed, macrophage targeted nanoparticles (19) may be an intriguing next step for targeted delivery of RvD2 to plaques.

Along these lines, targeting inflammation can lead to immunosuppression, which can become a huge problem, especially for progressive long-term diseases like atherosclerosis. RvD2 is not immunosuppressive in mice. Indeed, RvD2 limits polymicrobial sepsis by boosting phagocyte function. In the context of atherosclerosis, there are numerous reports that suggest boosting phagocyte function can limit plaque necrosis. Therefore, RvD2 offers a therapeutic approach that targets clearance and repair as opposed to simply blocking inflammation at its source.

## Non-standard abbreviations

GPR18: G-protein coupled receptor 18 (RvD2’s receptor)
SPMs: Specialized pro-resolving mediators
RvD2: Resolvin D2

## Author Contributions

M.L. performed all *in vivo* experiments and lesional analyses. S.W. contributed to the bone marrow transplant lesional analysis. M.L. contributed to *in vitro* experiments regarding the characterization of the fl/fl and mKO mice. H.A.M. performed the *in vitro* BMDM experiments. M.S. supervised selected experiments and contributed to the writing of the manuscript. G.F. conceived the overall design of the experiments, performed analysis on selected experiments and contributed to the writing of the manuscript.

